# Rationally designed protein bandpass filters for controlling cellular signaling with chemical inputs

**DOI:** 10.1101/2023.02.10.528077

**Authors:** Sailan Shui, Leo Scheller, Bruno E. Correia

## Abstract

Biological mechanisms that rely on signal integration and processing are fundamental for cell function. These types of capabilities are analogous to those found in electronic circuits where individual components perform operations on input signals. In electronics, bandpass filters are crucial components to narrow frequencies within a specified range and reject frequencies outside of that range. However, no generalizable protein-based components are currently available to mimic such processes in engineered biological systems, representing an unmet need in controllable modules. Here, we propose a rational design approach to create protein-based *chemically responsive bandpass filters* (CBP) which pass chemical concentrations within a range and reject concentrations outside of that range, showing an OFF-ON-OFF regulatory pattern. The CBPs were designed using structure-based approaches where we created a heterodimeric construct which the assembly is triggered by low concentration of a small-molecule, and this interaction is inhibited at high concentrations of the drug, effectively creating a bandpass filter. The CBPs have a multidomain architecture where we used known drug-receptors, a computationally designed protein binder and small-molecule inhibitors. Owing to the modularity of the system, each domain of the CBPs can be rationally fine-tuned to optimize its performance, including bandwidth, maximum response, cutoff concentration and fold changes. These CBPs were used to regulate cell surface receptor signaling pathways showing the capability to control cellular activities in engineered cells.

## Introduction

Cells are sophisticated signal processing units that respond and adapt to environmental and internal signals^1^. Synthetic biologists strive to engineer cells that process signals in a systematic, predictable, controllable, and integrated manner^2^. Chemically responsive protein modules are particularly useful tools for engineering artificial cellular activities controlled by external cues^1,3^. Currently available protein switches are designed to sense a specific chemical and control the activity of biological circuits through induced protein-protein association (e.g. rapamycin-induced FKBP:FRB dimerization^4^) or dissociation (chemically disruptable heterodimers^5^). Protein switches are capable to up- or down-regulate cellular activities, but require a high dilution of the chemicals to revert the controlled cellular activity from ON to OFF, or vice-versa^3^. However, cells perceive and respond to temporal and spatial stimulations in a timely and efficient manner^6^. For instance, processes that result in pattern formation are a hallmark of such coordinated and complex behavior. Current engineered protein components mostly uniformly up- or down-regulate outputs, and therefore engineered cellular systems often fail in replicating such sophisticated signal processing events.

The morphogen driven pattern organization is a hallmark example of coordination of cell behavior and signal detection of gradient concentrations. To mimic the morphogen dependent function, many efforts have been focused on developing band-detecting systems which analogizes the electronic bandpass filters^7^. An electronic bandpass filter can pass frequencies within a defined range and reject frequencies outside that range. Basu et al, engineered a bandpass filter by tuning a trans-repressor’s affinity for the acyl-homoserine lactone signaling molecule and dosing with different plasmids copies^8^. Similar work has also been demonstrated in bacteria, in which an externally tunable bandpass filter was built to tune enzymatic activities using IPTG^9,10^. Greber and his co-workers constructed a synthetic genetic network with bandpass characteristics in which output expression is only “on” across a window of low concentrations of tetracycline^11^. Bacterial transcription factors have also been designed to sense and respond to D-fucose and IPTG to mimic band-detecting behavior^12^. Other biological networks with band-detecting characteristics have been developed based on transcription factors, including detection of L-arabinose^13^, Nisin^14^, and Ampicilin^15^.

Despite inducible genetic circuities that have been constructed to perform band-detecting functions, there are no engineered protein-based sensors that can perform such behaviors. There are also many limitations in terms in choice of transcription factors and cell permeable small molecule inducers that can be used in genetic circuits. To overcome these limitations, a generalizable approach for engineering robust protein-based bandpass filters is needed. Hence, we rationally designed *Chemically responsive protein-based BandPass filters* (CBPs) that detect drug concentrations, pass concentrations within a certain range and reject concentrations beyond that range, resulting in an “OFF-ON-OFF” regulatory pattern. Each CBP consists of three distinct protein components: a designed binder, a drug_low_-receptor and a drughigh-receptor which are sensitive to drug disruption in low and high concentrations respectively. Each CBP undergoes an “ON-switch” process, where the designed binder dissociates from the drug_low_-receptor at low drug concentrations and consequently binds to the drug_high_-receptor, and an “OFF-switch” process, where the designed binder dissociates from the drug_high_-receptor at high drug concentrations (**Figure 1a**). Based on the unique behavior of CBPs, we identified five key parameters of CBPs, including fold changes between ON- and two OFF-states, maximum response, cutoff concentration, and bandwidth that can sustain the activity controlled by CBP at a defined level (**Figure 1b**). These protein-based CBPs are also highly modular, the binding affinities between drug-receptors and the designed binder as well as the drug sensitivities of drug-receptors can be modulated to adjust the parameters of the CBPs. Furthermore, the designed CBPs were incorporated in cell surface receptors to regulate signaling pathways externally and sensitively in a bandpass filtering fashion. Our results show the potential of rationally designed protein-based CBPs for diverse cellular applications.

**Figure 1:**
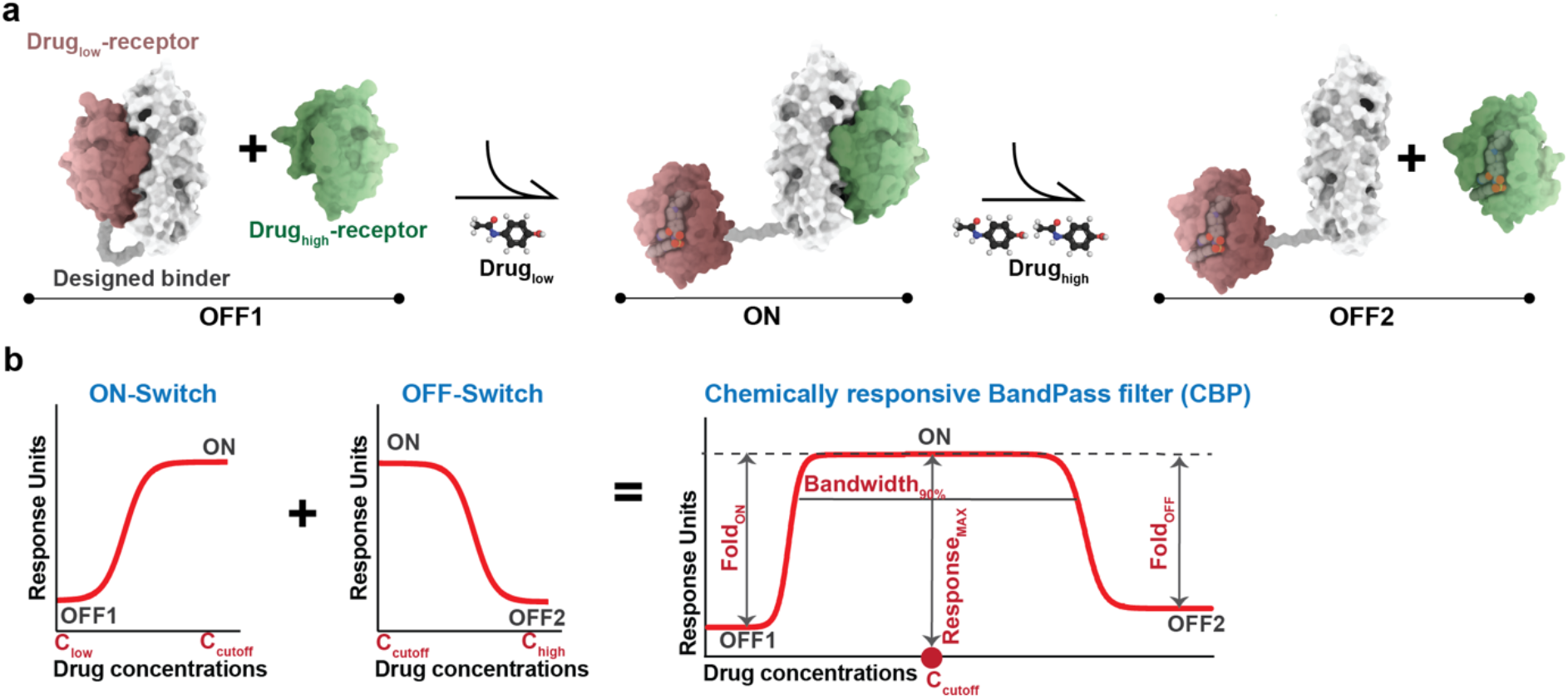
Design strategy and performance parameters of the chemically responsive protein-based bandpass filters (CBPs). **a)** Schematic representation of the CBP mode of action. Each CBP contains three proteins: the designed binder (white surface representation), and two drug-receptors (drug_low_ -receptor in red and drug_high_ -receptor in green). For low drug concentrations, CBPs switch to the ON-state, and in high drug concentration, CBPs switch to the OFF2 state. **b)** Design principles of the CBPs by integrating an ON-switch (from low drug concentration to cutoff drug concentration) and an OFF-switch (from cutoff drug concentration to high drug concentration), and the tunable parameters of each CBP, including cutoff concentration (C_cutoff_ = Drug concentration of Response_MAX_), bandwidth (Bandwidth_90%_ = Drug concentration range maintains >= 90% of Response_MAX_), fold changes (Fold_ON_ = Response_MAX_ /Response_OFF1_, Fold_OFF_ = Response_MAX_/Response_OFF2_) and the maximum response (Response_MAX_ = Maximum Response).

## Results

### Bandpass filter design by integrating ON- and OFF-switches

Analogous to electronic bandpass filters, we sought to design CBPs with an OFF-ON-OFF regulatory pattern depending on the concentration of a drug. To do so, three proteins will be used to construct a CBP: a designed binder that interacts with both drug-receptors, a drug receptor that is dissociated from the designed binder at low drug concentrations (drug_low_-receptor), and a drug receptor which can only be dissociated at high drug concentrations (drug_high_-receptor). The designed binder is fused to the drug_low_-receptor (OFF1 state, intramolecular dissociation), and is dissociated from the drug_low_-receptor to bind to the drug_high_-receptor at low drug concentrations (ON state, intermolecular association), and is dissociated from both drug-receptors at high drug concentrations (OFF2 state) (**Figure 1a**).

Biological activities controlled by CBPs are designed to be upregulated and then downregulated with increasing drug concentrations. We hypothesized that the up- and down-regulation could be mimicked by integrating a protein-based ON-switch, which turns biological activity ON from low to cutoff concentrations (from OFF1 state to ON state), and an OFF-switch which turns biological activity OFF from cut-off to high concentrations (from ON state to OFF2 state) (**Figure 1b, left**). Many parameters of CBPs can be characterized and modulated, beyond fold changes and maximum response, we also monitored the cut-off concentration indicating the highest response that can be achieved, and the bandwidth indicating the range of drug concentrations that retain the ON state (**Figure 1b, right**).

### A protein-based bandpass filter with dual drug control

We sought to construct a dual-drug controlled CBP analogue (dCBP) to test if the ON- and OFF-switching process can be combined in a protein-based architecture. To do so, we used orthogonal drug1-receptor and drug2-receptor which interact with the designed binder but respond specifically to their corresponding inhibitors. In such setting, the dCBP will turn ON in response to one drug and OFF in response to the other, thus mimicking bandpass filtering behavior (**Supplementary Figure 1**).

To do so, LD3 was used as the designed binder which has been previously designed to interact with BclXL and Bcl2 proteins^16^ as drug-receptors. The compound A-1155463 selectively interacts with BclXL with a KD lower than 10 pM, while its binding to Bcl2 is more than 7000-fold weaker^17^. In a surface plasmon resonance (SPR) drug competition assay, we confirmed that A-1155463 prevents BclXL from binding to LD3 with an IC_50_ of 106 nM, and does not affect the Bcl2:LD3 interaction with a detectable IC_50_ (**Supplementary Figure 2a**). This result shows that Bcl2 can be used as the drug2-receptor which is not affected by BclXL’s inhibitor. Thus, we constructed the first dCBP using Bcl2 and the fused BclXL-LD3 module, which is expected to be turned ON by A-1155463 and turned OFF by Venetoclax, referred to as dCBP_A11-Vene_. dCBP_A11-Vene_ was tested in the generalizable extracellular molecule sensor (GEMS) platform^18^ for regulating cell surface signaling pathways externally (**Figure 2a**). In the GEMS system, Bcl2 and BclXL-LD3 were fused to the engineered cytokine receptor chains. In this setting, dimerization of Bcl2 and BclXL-LD3 triggers JAK/STAT3 signaling, leading to the expression of a reporter protein (secreted alkaline phosphatase - SEAP). The dCBP_A11-Vene_ turned on reporter gene expression with a 100-fold increase in the presence of 100 nM A-1155463 and was effectively shut down by 1 µM Venetoclax (**Figure 2b**). The dual use of A-1155463 and Venetoclax did not show toxicity in reporter gene production (**Supplementary Figure 2b**) nor cell proliferation (**Supplementary Figure 2c**). The dCBP_A11-Vene_ showed a dose-dependent response with EC_50_ of 23 nM in the ON phase and IC_50_ of 11 nM in the OFF phase (**Figure 2c**).

**Figure 2:**
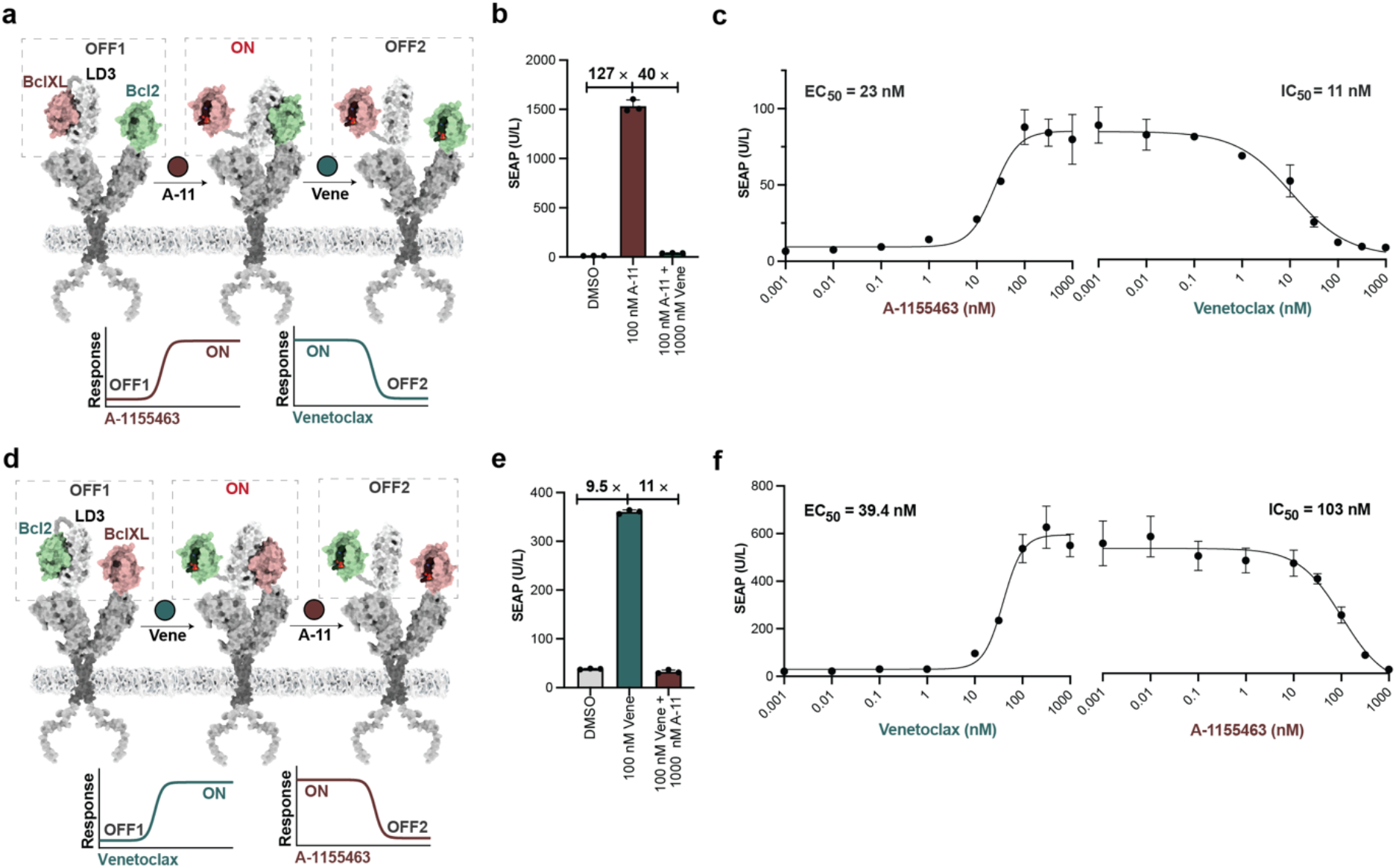
dCBPs controlled by two sequential chemical inputs mimicking the CBP behavior. **a)** Scheme of dCBP_A11-Vene_ architecture (Bcl2 & BclXL-LD3 in each receptor chain, switching between states for ON- and OFF-regulatory phases. The dCBP_A11-Vene_ is initially in the OFF1 state and can be switched to the ON state by the BclXL inhibitor A-1155463 (A-11; red circle) which can be switched to the OFF2 state by the Bcl2 inhibitor Venetoclax (Vene; green circle). The curves on the bottom show the idealized output target for switching between states. **b)** Fold-change of dCBP_A11-Vene_-GEMS acitivity between no drug treatment (DMSO) versus 100 nM A-1155463 (ON) versus 100 nM A-1155463 and 1 µM Venetoclax (OFF), showing SEAP expression after 24 h. Each bar represents the mean of three biological replicates ±s.d., overlaid with the original data points. **c)** Dose responses in engineered cells expressing dCBP_A11-Vene_ in the GEMS platform. Each data point represents the mean ±s.d. of three replicates and the EC_50_ s of ON-phase and IC_50_ s of the OFF-phase were calculated individually using four-parameter nonlinear regression. **d)** Scheme of dCBP_Vene-A11_ architecture (BclXL & Bcl2-LD3 in each receptor chain switching between ON- and OFF-phases. The dCBP_Vene-A11_ can be turned from the OFF1 state to the ON state by the Bcl2 inhibitor Venetoclax (Vene; green circle). It can be switched to OFF2 state with the BclXL inhibitor, A-1155463 (A-11; red circle). The curves on the bottom show the idealized output target for switching between states. **e)** Fold-change of dCBP_Vene-A11_-GEMS activity between no drug treatment (DMSO) versus 100 nM Venetoclax (ON) versus 100 nM Venetoclax and 1 µM A-1155463 (OFF), showing SEAP expression after 24 h. Each bar represents the mean of three biological replicates ±s.d., overlaid with the original data points. **f)** Dose responses in engineered cells expressing the dCBP_Vene-A11_ in the GEMS platform. Each data point represents the mean ±s.d. of three replicates and the EC_50_ s of ON-phase and IC_50_ s of the OFF-phase were calculated using four-parameter nonlinear regression.

Venetoclax selectively binds to Bcl2 rather than to BclXL with 500-fold stronger affinity^19^. Venetoclax also shows great selectivity in inhibiting LD3 interactions with Bcl2 (IC_50_ = 67 nM) while not affecting the interaction between BclXL and LD3 (**Supplementary Figure 2d**). Using the same principle as before, we constructed dCBP_Vene-A11_ which can be turned ON by Venetoclax and turned OFF by A-1155463 (**Figure 2d**). The dCBP_Vene-A11_ showed a close to 10-fold increase and decrease in its ON- and OFF-phases, respectively (**Figure 2e**). The dCBP_Vene-A11_ showed a dose-dependent response with and EC_50_ of 39 nM in the Venetoclax controlled ON phase and IC_50_ of 103 nM in the A-1155463 controlled OFF-phase (**Figure 2f**). The combined use of A-1155463 and Venetoclax on dCBP_Vene-A11_ did not cause cellular toxicity (**Supplementary Figure 2e-f**).

These results show that the dCBPs can regulate cell surface receptor signaling in an OFF-ON-OFF pattern in response to two drugs serving as a proof of concept for our design strategy.

### A chemical bandpass filter responsive to one chemical input

To test whether the dCBP strategy translates into a single chemical input CBP design, we tested the dCBP_A11-Vene_ and dCBP_Vene-A11_ against the Bcl2 family inhibitor Navitoclax, which binds to both BclXL and Bcl2 with different potencies^20^. We compared Navitoclax Bcl-XL:LD3 or Bcl2:LD3 complex disruption in the GEMS platform individually, which showed approximately 4.4-fold more potent activity to Bcl-XL than to Bcl2, IC50s of 403 nM and 1773 nM, respectively (**Supplementary Figure 3a-b**).

Next, we tested a CBP responsive to Navitoclax (CBP_Navi_) that dissociates LD3 from the fused BclXL-LD3 complex to interact with Bcl2 at low concentrations, and dissociates LD3 from Bcl2 at high concentrations (**Figure 3a**). CBP_Navi_ were used to control receptor activity in the GEMS platform, which can be maximally activated at a cutoff concentration of 320 nM with approximate 9-fold increase of reporter gene expression, and can be turned OFF at 10 µM (**Figure 3b)** that do not interfere with reporter gene expression nor cell proliferation (**Supplementary Figure 3c-d**). We also characterized the ON- and OFF-kinetics of CBP_Navi_, showing an EC_50_ of 65 nM in the ON phase and an IC_50_ of 2820 nM in the OFF-phase (**Figure 3c**).

**Figure 3:**
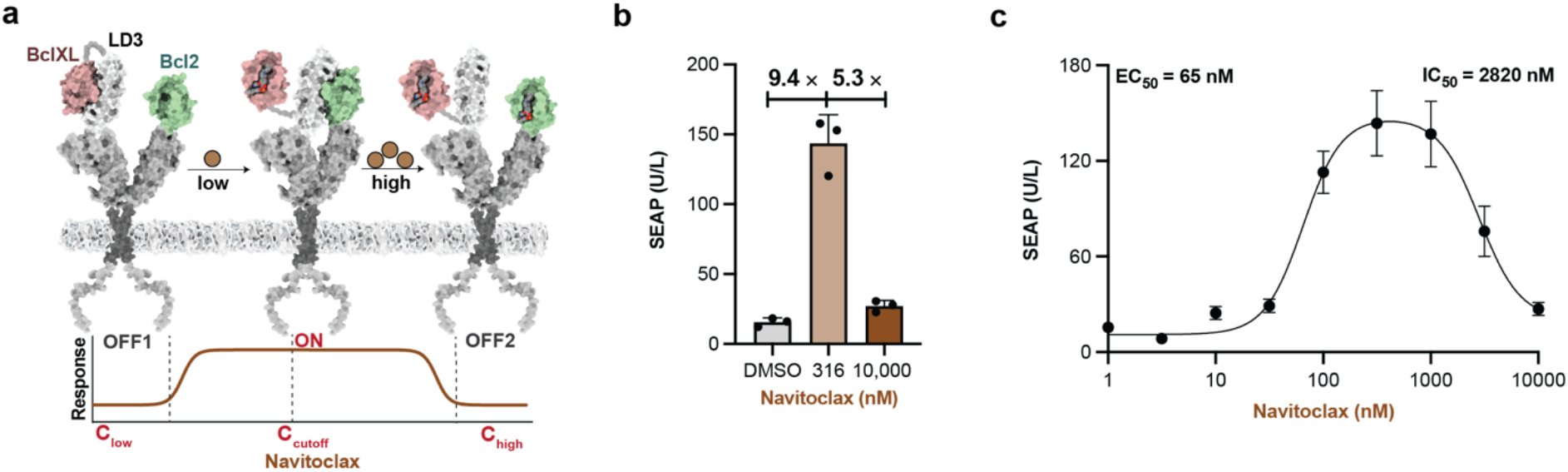
Schematic representation and characterization of CBP_Navi_. **a)** Scheme of CBP_Navi_ and its response to different drug concentrations. **b)** Fold-change in reporter gene expression between no drug treatment (DMSO; OFF1) versus 316 nM (ON) versus high concentration 10,000 nM (OFF2), 24 h after drug addition. Each bar represents the mean of three biological replicates ±s.d, overlaid with a scatter dot plot of the original data points. **c)** Dose-response to Navitoclax in engineered cells expressing CBP_Navi_. Each data point represents the mean ± s.d. of three biological replicates. EC_50_ of ON-phase and IC_50_ of the OFF-phase were calculated using four-parameter nonlinear regression.

The CBP_Navi_ was constructed based on the different drug sensitivities between BclXL and Bcl2 towards Navitoclax, implying that rational design of drug-receptors with different drug sensitivities can lead to a general strategy for the engineering of band-pass filter protein devices.

### Rational design of CBPs

To rationally design a CBP responding to a chemical input, we used BclXL as the drug_low_-receptor, LD3 as the designed binder, and a designed BclXL variant as drug_high_-receptor (referred to as BclXL_high_) to construct a CBP responsive to BclXL inhibitors (**Figure 4a**). To design BclXL_high_, we used a multi-state design computational protocol described elsewhere^16^ (**Details in method**) and generated six variants BclXL_high_ -v(1-6), which retain binding to LD3 but have higher resistance to A-1155463 (**Figure 4b, Supplementary Table 1**). We screened the candidate receptors in the CBP architecture and we observed that BclXL_high_ -v4-6 maintained strong resistance to A-1155463, while BclXL_high_-v1-3 showed a significant decrease when the drug concentration reached 1 µM (**Supplementary Figure 4a**). Thus, BclXL_high_-v1-3 were chosen as the functional drug_high_-receptor resulting in three A-1155463-controlled CBPs (referred to as CBP_A11_-v(1-3)) (**Figure 4b**). A-1155463 does not perturb reporter gene expression nor cell survival at the highest concentration of 10 µM (**Supplementary Figure 4c**,**d**).

**Figure 4:**
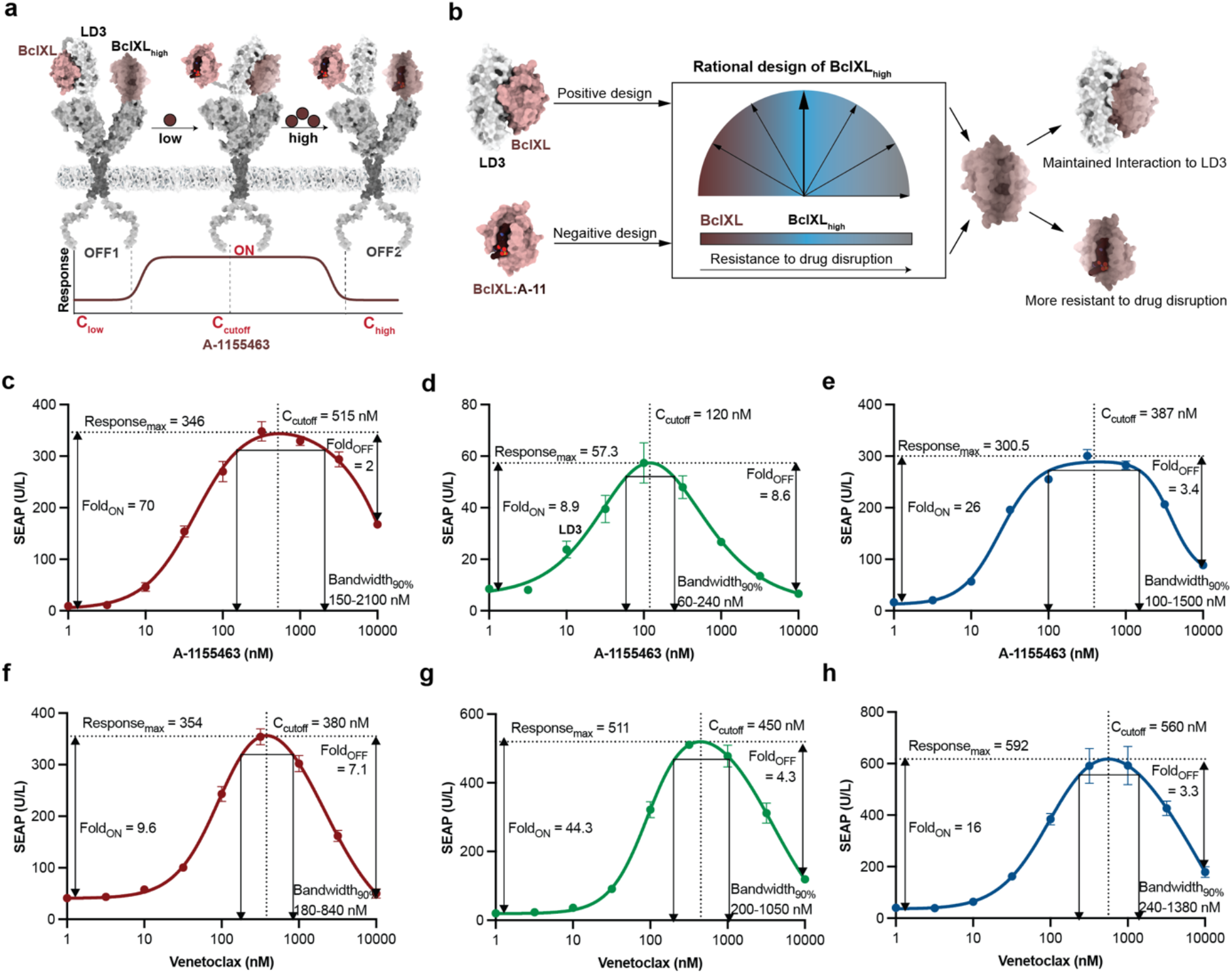
Rational design of CBPs that respond to small-molecule drugs. **a)** Scheme of CBP_A11_ and its response in different ranges of drug concentration. **b)** BclXL_high_ designs were generated to perform bandpass filtering functions in response to different concentrations of A-1155463. **c-e)** Dose responses of CBP_A11_-v1 (red), CBP_A11_-v2 (green), CBP_A11_-v3 (blue) in engineered cells. Each data point represents the mean ± s.d. of three replicates and the curves were calculated using Bell-shaped fitting. **f-h)** Dose responses of CBP_vene_-v1 (red), CBP_vene_-v2 (green), CBP_vene_-v3 (blue) in engineered cells. Each data point represents the mean ± s.d. of three replicates and the curves were calculated using Bell-shaped fitting.

All CBPA11-v(1-3) showed distinctive ON-phases and OFF-phases in response to different concentration ranges of A-1155463. CBP_A11_-v1 and CBP_A11_-v3 showed an ON-phase with a Fold_ON_ of 70-fold and 26-fold reporter expression over OFF1, respectively; however, they did not efficiently shut down in the OFF2-phase (**Figure 4c,e**). CBP_A11_-v2 demonstrated Fold_ON_ of 9-fold with 100 nM A-1155463 but shut down to background level at 10 µM (**Figure 4d**). We hypothesize that the fold changes (Fold_ON &_ Fold_OFF_) are related to the binding affinities between drug_high_-receptor and LD3 protein (BclXL_high_-v1: 44 nM & BclXL_high_ -v3: 37 nM vs BclXL_high_-v2: 159 nM) with the stronger binder contributing to the higher Fold_ON_ and lower Fold_OFF_ (**Supplementary Figure 5a**). The tighter interaction of BclXL_high_-v1 & v3:LD3, requires a higher drug concentration to dissociate the intermolecular interaction, resulting in a stronger activation and an ineffective disruption in OFF2-phases. Conversely, the weak interaction between BclXL_high_-v2 and LD3 resulted in a lower receptor activation but complete disruption in the OFF-phase of CBP_A11_-v2 (**Figure 4d)**.

Next, we characterized the C_cutoff_ and Bandwidth_90%_ parameters of CBP_A11_-v(1-3). CBP_A11_-v2 showed the lowest C_cutoff_ at around 120 nM, and CBP_A11_-v1&v3 both have a C_cutoff_ above 300 nM (**Figure 4c-e**). CBP_A11_-v2 has the narrowest Bandwidth_90%_ of 80-240 nM. In contrast, CBP_A11_-v1 can has a Bandwidth_90%_ of 150 nM to 2100 nM (**Figure 4c-e**). This data shows that the drug sensitivity of BclXL_high_ -v(1-3) to A-1155463 (**Supplementary Figure 5b)** is well translated into the differences of their C_cutoff_ and Bandwidth_90%_ of CBP_A11_-v(1-3). The more sensitive BclXL_high_ comparing to wildetype BclXL, the lower C_cutoff &_ Response_MAX_, and the narrower the Bandwidth_90%_. For instance, BclXL_high_-v1 has the strongest drug resistance to A-1155463 which showed a correspondingly highest C_cutoff_ and the widest Bandwidth_90%_.

Next, we constructed a Venetoclax (FDA approved Bcl2 inhibitor) responsive CBP (CBP_vene_) using Bcl2, LD3 and a designed drug_high_-receptor based on Bcl2 (Bcl2_high_) (**Supplementary Figure 6a)**. Using multi-state design, we generated eight Bcl2_high_ variants (referred to as Bcl2_high_-v(1-8)) which were designed for lower affinity to Venetoclax compared to native Bcl2 (**Supplementary Figure 6b, Supplementary Table 2**). CBP constructs including Bcl2_high_-v(1-8) were screened and Bcl2_high_-v2, Bcl2_high_-v4, Bcl2_high_-v5 presented the desired behavior with a sharp decrease when Venetoclax exceeds 100 nM (**Supplementary Figure 6c**). We confirmed that 10 µM Venetoclax is not toxic for reporter gene expression and cell proliferation (**Supplementary Figure 6d-e**).

CBP_vene_-v2 showed a Fold_ON_ and a Fold_OFF_ of about 9-fold (**Figure 4d**). CBP_vene_-v4 has a higher Fold_ON_ (44-fold) but does not completely shut off at 10 µM with a Fold_OFF_ of 3.4 (**Figure 4e**). CBP_vene_-v5 displayed a Fold_ON_ of 16-fold, the highest C_cutoff_ at 560 nM and the widest Bandwidth_90%_ with a range of 240-1380 nM (**Figure 4f**). Notably, the binding affinities between Bcl2_high_-v(2,4,5) and LD3 are relatively close (**Supplementary Figure 7a**), indicating that their respective drug resistance is critical for regulating the C_cutoff_ and Bandwidth_90%_. For instance, the Bcl2_high_-v4 is more resistant than Bcl2_high_-v2 (**Supplementary Figure 7b**) leading to the higher C_cutoff_ (CBP_vene_-v4: 450 nM vs CBP_vene_-v2:380 nM) and the wider Bandwidth_90%_ (CBP_vene_-v4: 200-1050 nM vs CBP_vene_-v2: 180-840 nM).

We developed CBPs controlled by BclXL and Bcl2 inhibitors by rationally designing drug_high_-receptors with reduced drug sensitivities and using in a multidomain architecture that can provide the output analogous to bandpass filters.

### Rational tuning of CBPs towards lower drug demand and larger bandwidth

We observed that CBP_A11_-v1 & CBP_A11_-v3 or CBP_vene_-v4 & CBP_vene_-v5 cannot be fully turned off at the highest drug concentration of 10 µM. However, very high drug concentrations may cause unwanted side effects due to toxicity. To improve the performance of these CBPs, we decided to rationally tune the binding affinities between LD3 and two drug-receptors in order to lower their C_cutoff_ and increase the shutdown efficiency.

Previously, we showed that mutations in the binding region of LD3 weaken the interaction of BclXL:LD3, thus lowering the drug demand to dissociate BclXL:LD3^16^ (**Figure 5a**). We expect to obtain the same effect of such mutations in the context of the CBPs, ultimately requiring lower drug concentrations to turn ON and shut OFF the CBPs. We replaced LD3 in CBP_A11_-v(1-3) with LD3-v(1-3) to test if they respond to lower concentrations of A-1155463. LD3-v3 based designs successfully shifted the C_cutoff_ of all three CBP_A11_-v(1-3) to lower drug concentrations (**Supplementary Table 3, Supplementary Figure 8a-c**). CBP_A11_-v1_LD3-v3_ and CBP_A11_-v3_LD3-v3_ were the best performers in terms of fold change and both shifted their C_cutoff_ from above 500 nM to below 100 nM and effectively shut down at 10 µM (**Figure 5b**). However, the new LD3-v3 based CBPs also showed a lower level of ON response **(Supplementary Figure 8 d-e)**.

**Figure 5:**
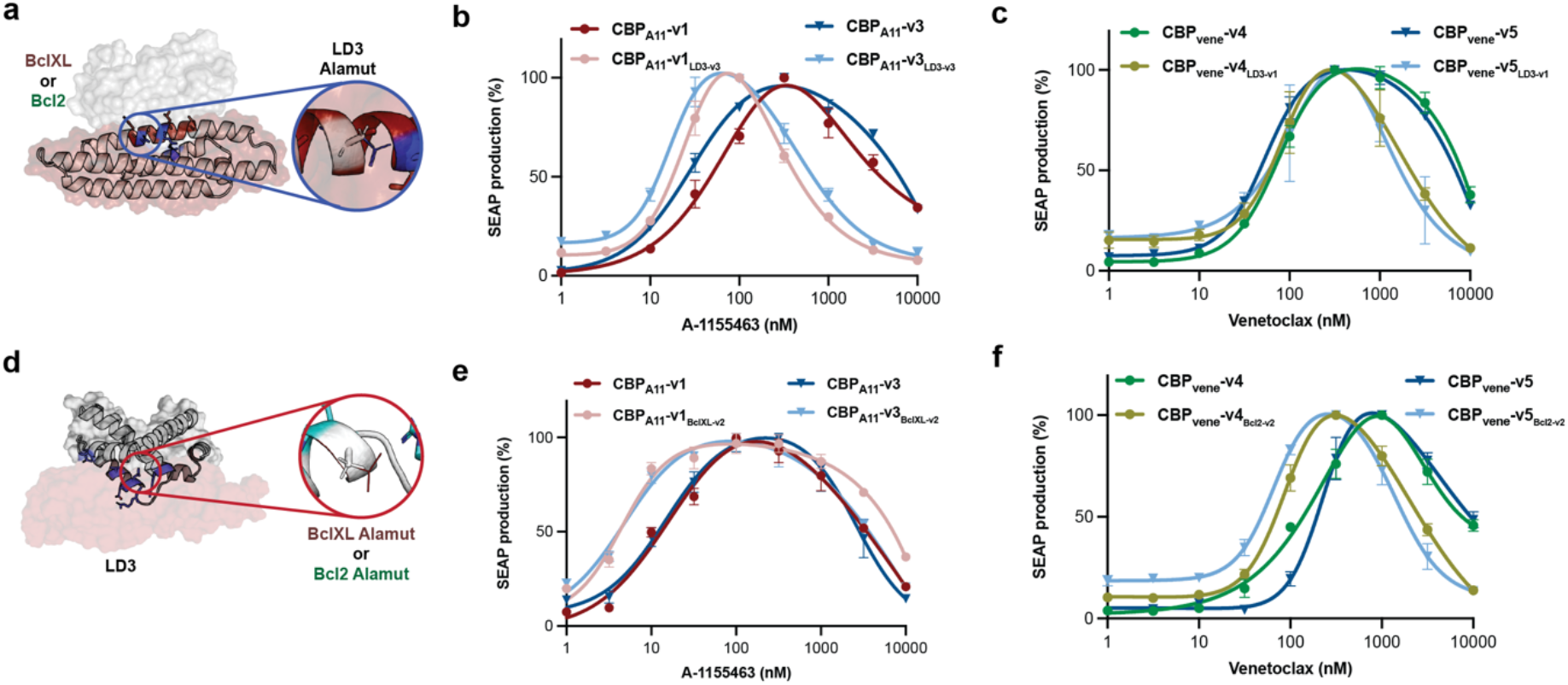
Rational tuning of binding affinities between the LD3 binder and drug-receptors can further modulate the characteristics of CBPs. **a)** Rational tuning of binding affinities between LD3 (in red) and the drug-receptor (BclXL or Bcl2 in white surface). Positions were selected and mutated to alanine on LD3 to decrease the binding affinity. **b)** Normalized dose-responses of CBP_A11_-v1 with and without LD3-V3. **c)** Normalized dose-responses of CBP_vene_-v4 and CBP_vene_-v5_LD3-v1_ with and without LD3-v1. **d)** Scheme of the rational tuning of binding affinities between LD3 and the drug-receptor by alanine scanning on BclXL or Bcl2 proteins. **e)** Normalized drug dose-dependent responses of CBP_A11_-v1 (red) and CBP_A11_-v3 (blue) compared with CBP_A11_-v1_BclXL-v2_ (pink) and CBP_A11_-v3_BclXL-v2_ (turquoise) in engineered cells. **f)** Normalized drug dose-dependent responses of CBP_vene_-v4 (green) and CBP_vene_-v5 (blue) compared with CBP_vene_-v4_Bcl2-v2_ (asparagus) and CBP_vene_-v5_Bcl2-v2_ (turquoise) in engineered cells. **b**,**c**,**e**,**f)** Each data point represents the mean ±s.d. of three replicates and the curves were computed using Bell-shaped fitting.

We also replaced LD3 with LD3-v(1-3) in CBP_vene_-v(2,4,5) to test whether they respond to lower concentration of Venetoclax. Unlike CBP_A11_-v(1-3), LD3-v3 replacement in CBP_vene_-v(2,4,5) showed leaky background activities that failed to maintain the bandpass filtering behavior (**Supplementary Figure 9a-c**). The LD3-v1 replacement shifted both CBP_vene_-v4_LD3-v1_ and CBP_vene_-v5_LD3-v1_ to lower drug concentration ranges with no significant decrease in C_cutoff_s compared to CBP_vene_-v4 or CBP_vene_-v5 respectively (**Figure 5c, Supplementary Figure 9d-e**). We found that the binding affinity of LD3-v3:BclXL complex in CBP_A11_-v1_LD3-v3_ and CBP_A11_-v3_LD3-v3_ is 5000-fold weaker compare to the wildtype LD3:BclXL complex^16^, which can drastically lower the C_cutoff_ of CBP_A11_-v1 and v3. However, the binding affinity of LD3-v1:Bcl2 in CBP_vene_-v4_LD3-v1_ and CBP_vene_-v5_LD3-v1_ is less than 200-fold weaker than LD3:Bcl2^16^, which was not capable of shifting the C_cutoff_s significantly. We confirmed that replacement of LD3 with weaker variants was able to adjust the response of CBP_A11_ and CBP_Vene_ to lower drug concentrations, and that only a significant decrease in the binding affinity of LD3 to its variants was able to shift C_cutoff_s noticeably.

Next, we tested if we could rationally engineer the drug_low_-receptor to increase the sensitivity of the OFF1 to ON regulatory phase (**Figure 5d**). Since the switch from ON to OFF2 should not be affected, we hypothesized that this strategy could also increase bandwidth. We scanned residues at the interface of BclXL:LD3 complex, selected three residues on BclXL and mutated them individually to alanine. We selected them to affect the binding affinity in mild, moderate and strong levels (**Details in methods, Supplementary Table 4**). Among the BclXL variants (referred to as BclXL-v(1-3)), BclXL-v2 emerged as the best candidate since BclXL-v1 and v3 failed to function as bandpass filters in a CBPA11 setting (**Supplementary Figure 10a-c**). We found that both CBP_A11_-v1_BclXL-v2_ and CBP_A11_-v3_BclXL-v2_ showed similar C_cutoff_s and a wider bandwidth compared to CBP_A11_-v1 and CBP_A11_-v1, respectively (**Figure 5e)**. While OFF-phase kinetics remained similar, CBP_A11_-v1_BclXL-v2_ and CBP_A11_-v3_BclXL-v2_ turned ON at lower drug concentrations than CBP_A11_-v1 and CBP_A11_-v3 reflecting the weaker interaction of BclXL-v2:LD3 than BclXL:LD3 (**Supplementary Figure 10d-e**).

Applying the same principle to CBP_vene_-v(2,4,5), we also introduced alanine mutations into Bcl2 (**Details in method, Supplementary Table 5**), referred to as Bcl2-v(1-6). Based on the screening results (**Supplementary Figure 11a-c**), Bcl2-v2 was used for constructing CBP_vene_-v4_Bcl2v2_ and CBP_vene_-v5_Bcl2v2_. Both CBP_vene_-v4_Bcl2v2_ and CBP_vene_-v5_Bcl2v2_ demonstrated a lower C_cutoff_ and unexpectedly also shut down in lower concentrations comparing to CBP_vene_-v4 and CBP_vene_-v5, respectively (**Figure 5f, Supplementary Figure 11d-e**).

Altogether, we showed the improved performance of CBP_A11_ and CBP_vene_ with better Fold_OFF_s and lower C_cutoff_s by rationally tuning the binding affinities between LD3 and two drug-receptors. We also re-designed the drug_low_-receptors of CBP_A11_ and CBP_vene_, BclXL and Bcl2 respectively, which shifted the ON-regulatory phase to lower drug concentration ranges to tune bandwidth and C_cutoff_s of CBPs.

## Discussion

In contrast to electrical devices, cells as biological computing units use chemical inputs and biological molecules for signal processing. Protein switches have been designed to turn cellular activity ON or OFF, however, protein modules that perform bandpass filter behavior independent of engineered transcription factors remain unexplored. Here, we use computational protein design to develop protein-based bandpass filters with tunable behavior that detect drug concentrations and regulate cellular activities in an OFF-ON-OFF pattern.

First, we mimicked the behavior of bandpass filters by combining two drugs as ON- and OFF-signals. This approach has been attempted using engineered bacterial transcription factors to react to two chemicals^12^, however, it lacks transferability when different transcription factors or controlled genetic circuits are required. We presented here a novel strategy to engineer protein-based bandpass filters by rationally designing a drug_high_-receptor that remains bound to the designed binder (ON) and can only be dissociated at high drug concentrations (OFF). In this way, each CBP responds to a single chemical input and sense the ON- and OFF-signals dependent on the chemical concentrations. In total, we designed 10 CBPs based on BclXL and Bcl2 inhibitors producing different cutoff concentrations from below 100 nM (e.g. CBP_A11_-v1_LD3-v3_) to above 500 nM (e.g. CBP_vene_-v5), as well as regulatory bandwidths with different ranges (e.g. 60-240nM of CBP_A11_-v2 versus 100-1500 nM of CBP_A11_-v3). By rationally mutating the designed binder (LD3), CBPs can be tuned to perform with higher drug sensitivity in the trade-off with maximal response. We also demonstrated that by modulating different drug sensitivities of drug_low_-receptors, the greater the difference in drug sensitivity between drug_high_-receptor and drug_low_-receptor, the greater the bandwidth. These results demonstrate that rational protein design can be used to design and tune CBPs.

Protein-based CBPs function by induced protein-protein interactions, which can be modular and less restricted than intracellular genetic circuit based bandpass filters^21^. While previous synthetic bandpass filter systems sensed small molecules and produce a transcriptional response^8,11,12^, they did not enable the control of other cell functions such as signaling pathways. We utilized the designed CBPs to control engineered cytokine receptors which detect extracellular signals and regulate intracellular JAK/STAT3 signaling pathways. This application demonstrated the potential of CBPs in receptor engineering for sensing external cues, which is a first step towards sensing peptide or protein targets with bandpass behavior. Such systems could ultimately lead to the integration of bandpass filters into engineered cell communication networks for synthetic morphology or other autonomously controlled cell consortia^22,23^.

CBP-regulated systems have defined ranges of drug concentrations to sustain specific activity levels, in contrast to the simpler ON/OFF protein switch-based systems. For instance, the CBP_A11_-v1 can sustain 90% of receptor activation within drug concentrations from 150 nM to 2.1 µM. As a possible safety feature, exceeding the cutoff concentration, could serve as negative feedback to tune down the output in potential therapeutic applications. This drug concentration-dependent pattern of CBPs can be applied in controlling biological or therapeutical activities which require self-regulatory feedbacks from their inputs. We envision that the CBPs will be valuable components to control signal processing and to engineer sophisticated cellular functions.

## Methods and Materials

### Multistate design of single drug reversible Bcl-XL and Bcl2

As reported previously, a set of residues in the receptor protein’s (BclXL/Bcl2) binding site was selected for redesign^16^. From this set, a number of mutations were evaluated for binding energy to the binder protein (LD3) (positive design) or the drug (negative design). Afterwards all mutations were ranked according to the difference in energy between the positive design and negative design. The structure of BclXL bound to A-1155463 (PDB: 4QVX) was used for the negative design strategy, while the model of BclXL bound to LD3 (based on the BclXL:BIM BH3 structure with PDB: 3FDL) was used for positive design. Six BclXL residues in the binding site of A-1155463 (E98, R102, F105, T109, S145 and A149) were manually selected for redesign due to their proximity to drug moieties and relative distance to LD3 in the positive design structure. Each of these residues was allowed to mutate to residues with similar size/properties, restricted to a maximum of two simultaneous mutations from wildtype: E98: {E/S}, R102: {F/R/K/D/E/H}, F105:{F/L/V/I/A}; T109: {S/A/T/L/V}; S145: {S/D/E/V/A}; A149:{V/A/L/I}. Six sequences BclXLhigh-v1 (T109L, A149L), BclXLhigh-v2 (A149V), BclXLhigh-v3 (R102F, T109V), BclXLhigh-v4 (E98S,R102E, F105I), BclXLhigh-v5 (R102E, F105I, T109L), and BclXLhigh-v6 (E98S,R102E, F105I, T109L) were generated to test for single drug reversible BclXL. All sequences are listed in Supplementary Table 1.

The structure of Bcl2 bound to Venetoclax (PDB: 6O0K) was used for the negative design strategy, while the model of Bcl2 bound to LD3 (PDB: 6IWB) was used for positive design. Five Bcl2 residues in the binding site of Venetoclax (A100, D103, V148, V156 and Y202) were manually selected for redesign due to their closeness to drug moieties and relative distance to LD3 in the positive design structure. Each of these residues was allowed to mutate to amino acids with similar size/properties, restricted to a maximum of two simultaneous mutations from wildtype: A100: {A/S/T/V}, D103: {D/N/E/Q/S}, V148: {V/I/L/M/T}, V156: {V/I/L/M/T} and Y202: {Y/W/F/H/R/K/Q/E}. Three sequences srBcl2-v1(V156I, Y202H), srBcl2-v2(D103N, Y202H) and srBcl2-v3(A100T_D103S) were selected from the top results. Five sequences were generated based on the observation on Venetoclax resistance in clinical trials where G101V occurs alone or together with D103Y and D103E. Hence, we constructed srBcl2-v4(G101V), srBcl2-v5(D103Y) and srBcl2-v6(G101V,D103Y), srBcl2-v7(D103E), srBcl2-v8(G101V,D103E). All sequences are listed in Supplementary Table 2.

### AlaScan on LD3 and drug-receptor proteins

LD3-v(1-3) variants were designed previously^16^, sequences of LD3-v(1-3) are listed in Supplementary Table 3. The BclXL protein in cmplex with LD3 was computationally redesigned for a range of decreasing binding affinities. Rosetta’s alanine scanning filter was used to evaluate the change in ΔΔG for the LD3:BclXL complex (PDB id: 3FDL) upon mutating each of the 22 residues in the interface of BclXL to Alanine. The resulting list was then sorted by the change in ΔΔG, and three residues with positive levels of change in ΔΔG were selected: Q90 (0.33 REU), L90 (1.17 REU) and R99 (4.78 REU), where higher REU values are predicted to result in greater affinity losses. The three mutations were selected to provide a ‘gradient’ of affinities between LD3 and BclXL, sequences of BclXL-v(1-3) are listed in Supplementary Table 4. By the same principle, Rosetta’s alanine scanning filter was used to evaluate the change in ΔΔG for the LD3:Bcl2 complex (PDB id: 6IWB) upon mutating each of the 32 residues in the interface of BclXL to Alanine. The resulting list was then sorted by the change in ΔΔG, and six residues with positive levels of change in ΔΔG were selected: F63 (6.29 REU), V92 (1.29 REU), E95 (1.50 REU), L96 (0.88 REU), R105 (1.52 REU) and E111 (0.59 REU), where higher REU values are predicted to result in greater affinity losses. The six mutations were selected to provide a ‘gradient’ of affinities between LD3 and Bcl2, sequences of Bcl2-v(1-6) are listed in Supplementary Table 5.

### SPR assay for protein-protein binding affinities

Surface plasmon resonance (SPR) measurements were performed on a Biacore 8 K device (GE Healthcare). The mutual designed binder (LD3) was immobilized on a CM5 chip (GE Life Science) as a ligand with the concentration at 5 μg/ml for 120 seconds contact time in pH 4.5 sodium acetate solutions. Serial dilutions of the analytes (BclXL or Bcl2 and their variants) in HBS-EP buffer (10 mM HEPES, 150 mM NaCl, 3 mM EDTA and 0.005% Surfactant P20; GE Life Science) were flown over the immobilized chips. After each injection cycle, surface regeneration was performed using 10 mM NaOH (pH 11.95). Affinities (Kd) were obtained using a steady binding model of equilibrium model with Biacore 8 K evaluation software.

### SPR drug competition assay

Drug IC_50_s of disrupting heterodimers were measured on a Biacore 8K device. 5 µM of Analyte was mixed with 10 µM of BclXL’s or Bcl2’s inhibitors according to the analyte. The mixtures of analyte and drug were injected on the LD3 immobilized channel.

### Compounds

Venetoclax (>99.9%, Chemietek CT-A199), A1155463 (99.5%, Chemietek CT-A115), and NVP-CGM097 (100% optically pure, Chemitek CT-CGM097), were directly used without further purification. Venetoclax, A1155463, and NVP-CGM097 were each dissolved in DMSO as 10 mM stocks. Stocks were aliquoted and stored at -20 °C until use.

### Cell transfection and drug treatment

HEK293T cells were maintained in DMEM medium (Thermo Fisher) with 10% FBS (Gibco) and Pen/Strep (Thermo Fisher) at 37°C and 5% CO_2_ in a humidified incubator. Cells were maintained and split every two days at around 80% confluence.

HEK293T cells were seeded in 96-well plate 24 hours before transfection. The transfection mix in each well consisted of 3ng of constitutive human CMV promoter driven mammalian STAT3 expression vector (PhCMV-STAT3-pA), 30ng of STAT3-induced reporter expression (O_Stat3_-P_hCMVmin_-SEAP-pA), and 50ng of expression vectors for each GEMS receptor chain (PSV40-IgK-(drug_high_-receptor)-EpoRm-IL-6RBm-pA, and PSV40-IgK-(drug_low_-receptor-GGGGSX3-LD3)-EpoRm-IL-6RBm-pA). For the specific drug_high_-receptor and drug_low_-receptor used in different cellular assay, they are indicated as wildtype or with mutations. Then, plasmid DNA mixed with 50 μL opti-MEM (Thermo Fisher) and 600 ng of polyethyleneimine (24765-1, Polysciences, Inc.). For drug treatment experiments, drugs were added 12 hours post-transfection and incubated with cells for 24 hours before the SEAP reporter detection assay.

### Reporter detection assay

Secreted alkaline phosphatase (SEAP) activity (U/L) in cell culture supernatants was quantified by kinetic measurements at 405 nm (1 minute/measurement for 30 cycles) of absorbance increase due to phosphatase-mediated hydrolysis of para-nitrophenyl phosphate (pNPP). 4 - 80 µL of supernatant was adjusted with water to a final volume of 80 µL, heat-inactivated (30 min at 65 °C), and mixed in a 96-well plate with 100 μL of 2 × SEAP buffer (20 mM homoarginine (FluorochemChemie), 1 mM MgCl2, 21% (v/v) diethanolamine (Sigma Aldrich, D8885), pH 9.8) and 20 μL of substrate solution containing 20 mM pNPP (Sigma Aldrich, 71768).

### Statistics

Binding affinities of SPR drug competition assays were calculated using three-parameters nonlinear regression in GraphPad Prism (Version 8.3.0). Representative data of cell assays are presented as individual values and mean values (bars). n = 3 refers to biological replicates. All IC_50_ or EC_50_ values of cell assays reported were calculated using four-parameter nonlinear regression ± s.d. Bandpass features of cutoff concentrations and Bandwidth_90%_ were estimated based on the curve fitted using Bell-shaped curve non-linear regression in GraphPad Prism (Version 8.3.0).

## Contributions

SS and BEC conceived the work. SS performed the experimental characterization. SS, LS and BEC designed the experiments and analyzed the results. SS, LS and BEC wrote the manuscript.

### Data availability

The data supporting the findings of this study are available within the article and its Supplementary Information. Other data and reagents are available from the corresponding authors upon reasonable request.

## Acknowledgements

The computational simulations were facilitated by SCITAS at EPFL and by the Swiss National Supercomputer Center.

