## Supplementary material for "Rationally designed protein bandpass filters for controlling cellular signaling with chemical inputs": supp material

### Supplementary Figures

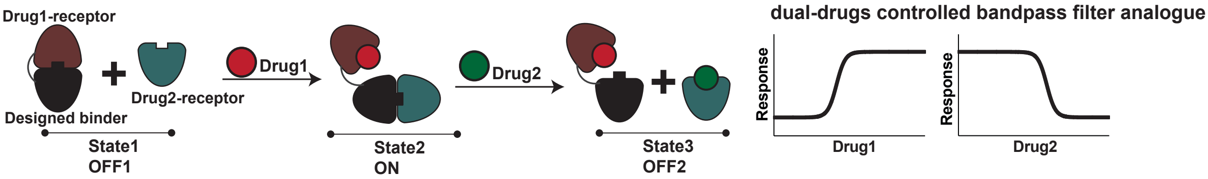


**Supplementary Figure 1: Architecture and mechanism of dCBP to produce bandpass filter behavior.**

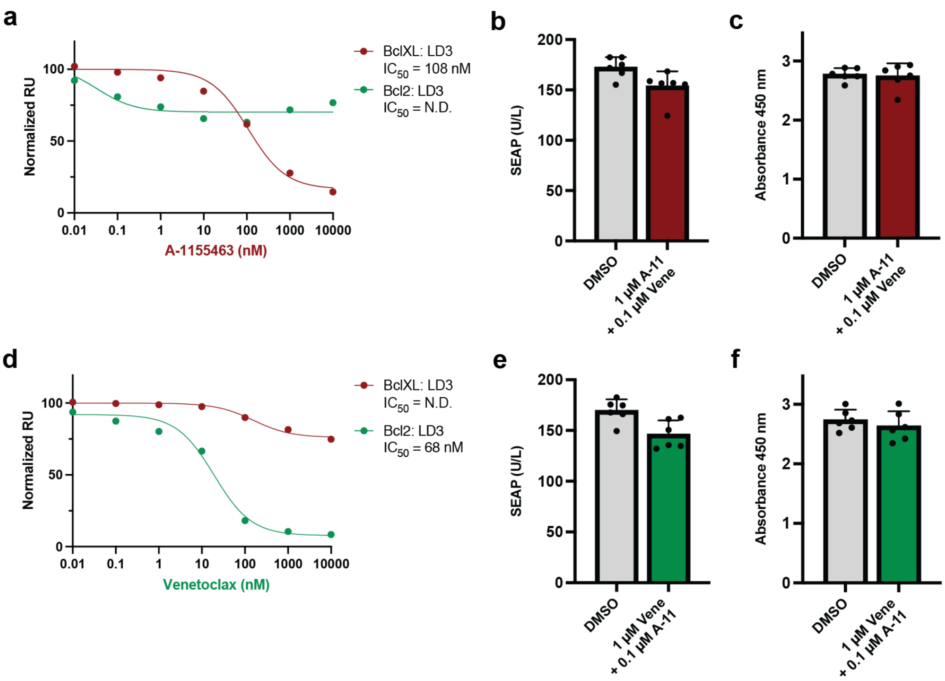


**Supplementary Figure 2: Bcl2 and BclXL selective drugs can be used for constructing dCBPs.** **a)** Apparent $IC_{50}$s for A-1155463 for dissociation of BclXL:LD3 and Bcl2:LD3 determined by SPR drug competition assay. **b,c)** Drug effects on cell viability and protein production of HEK293T cells. HEK293T cells were transfected with a plasmid for constitutive SEAP production and incubated with indicated drugs for 24 hours. SEAP quatification were performed to measure the protein production and WST-8 was used to test the cell proliferation. For cell proliferation assay, WST-8 was added after 24 hours of drug incubation, and measured its absorbance at 450 nm 4 hours later. Each bar represents the mean of six biological replicates ± s.d., overlaid with a scatter dot plot of the original data points. **d)** Apparent $IC_{50}$s for Venetoclax induce dissociation of BclXL:LD3 and Bcl2:LD3 determined by SPR drug competition assay. **e,f)** Drug effects on cell viability and protein production of HEK293T cells. HEK293T cells were transfected with a plasmid for constitutive SEAP production and incubated with indicated drugs for 24 hours. SEAP quatification were performed to measure the protein production and WST-8 was used to test the cell proliferation. For cell proliferation assay, WST-8 was added after 24 hours of drug incubation, and measured its absorbance at 450 nm 4 hours later. Each bar represents the mean of six biological replicates ± s.d., overlaid with a scatter dot plot of the original data points.

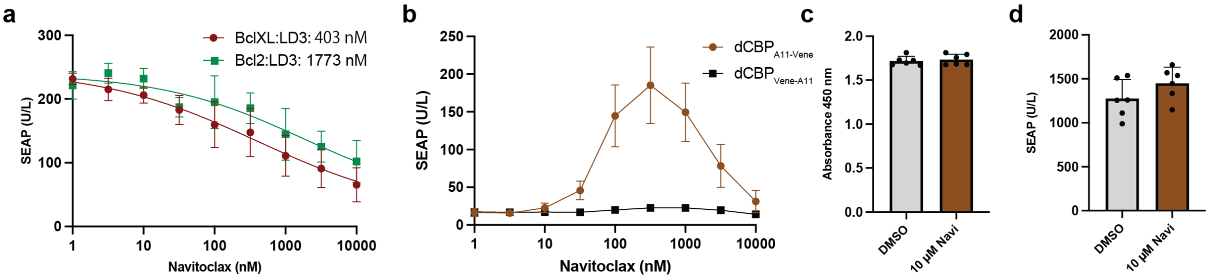


**Supplementary Figure 3: Navitoclax used to construct CBP$_{Navi}$.**

**a)** Dose responses of cells expressing GEMS receptors with BclXL:LD3 (PSV40-IgK-BclXL-EpoRm-IL-6RBm-pA and PSV40-IgK-LD3-EpoRm-IL-6RBm-pA) and Bcl2:LD3 (PSV40-IgK-Bcl2-EpoRm-IL-6RBm-pA and PSV40-IgK-LD3-EpoRm-IL-6RBm-pA) complexes disrupting by Navitoclax. **b)** Dose responses to Navitoclax of cells expressing the dCBP$_{A11\text{-}Vene}$ and dCBP$_{Vene\text{-}A11}$ in GEMS. Each data point represents the mean ± s.d. of three replicates. **c-d)** Navitoclax effects on cell viability and protein production of HEK293T cells. HEK293T cells were transfected with a plasmid for constitutive SEAP production and incubated with indicated drugs for 24 hours. SEAP quatification were performed to measure the protein production and WST-8 was used to test the cell proliferation. For cell proliferation assay, WST-8 was added after 24 hours of drug incubation, and measured its absorbance at 450 nm 4 hours later. Each bar represents the mean of six biological replicates ± s.d., overlaid with a scatter dot plot of the original data points.

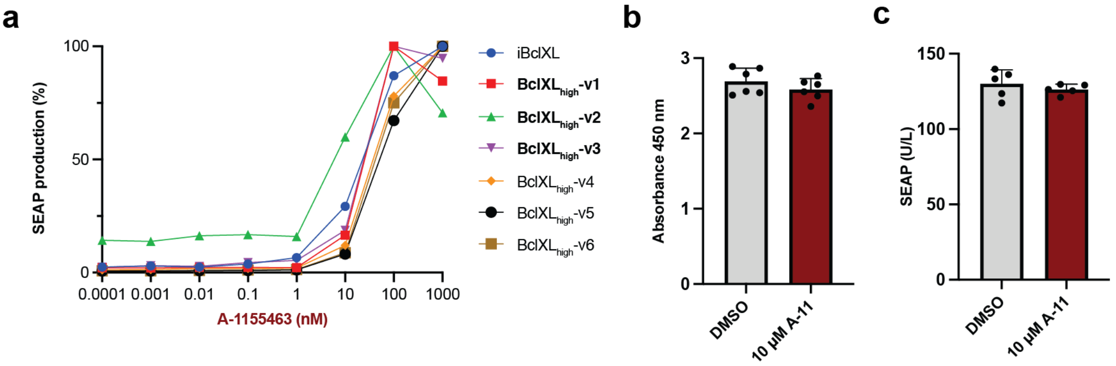


**Supplementary Figure 4: Design, screening and validation of BclXL$_{high}$.**

**a)** Screening of BclXL$_{high}$ variants in the GEMS platform. The designed binder LD3 was fused to the drug$_{low}$-receptor (PSV40-IgK-BclXL-GGGGSX3-LD3-EpoRm-IL-6RBm-pA) in one EpoR chain and BclXL$_{high}$ variants with the insensitive BclXL (iBclXL) as control in the other EpoR chain (PSV40-IgK-BclXL$_{high}$-EpoRm-IL-6RBm-pA). HEK293T cells were transfected with indicated plasmids, A-1155463 drug ranging from 1pM to 1 μM were added 12 hours post-transfection, then SEAP was measured 24 hours after drug treatment. **b-c)** A-1155463 effects on cell viability and protein production of HEK293T cells. HEK293T cells were transfected with a plasmid for constitutive SEAP production and incubated with indicated drugs for 24 hours. SEAP quatification were performed to measure the protein production and WST-8 was used to test the cell proliferation. For cell proliferation assay, WST-8 was added after 24 hours of drug incubation, and measured its absorbance at 450 nm 4 hours later. Each bar represents the mean of six biological replicates ± s.d., overlaid with a scatter dot plot of the original data points.

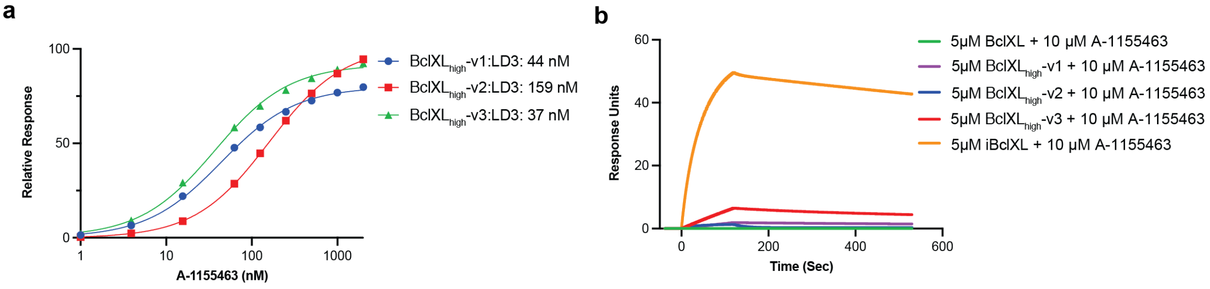


**Supplementary Figure 5: Binding affinities and drug resistances of BclXL$_{high}$-v(1-3).**

**a)** Dissociation constant measurement of BclXL$_{high}$-v(1-3) and LD3. Equilibrium curves are fitted using three-parameter non-linear regression. **b)** Comparison of A-1155463 sensitivity of BclXL$_{high}$-v(1-3) measured by SPR. 5 µM of BclXL or indicated variants were mixed with 10 µM of A-1155463 and injected over the LD3 immobilized chip to analyse the binding response. iBclXL remained stably bound to LD3 (orange), and BclXL was completely inhibited (green). BclXL$_{high}$-v1 (purple), BclXL$_{high}$-v2 (blue), BclXL$_{high}$-v3 (red) showed intermediate binding responses in the presence of 10 µM A-1155463.

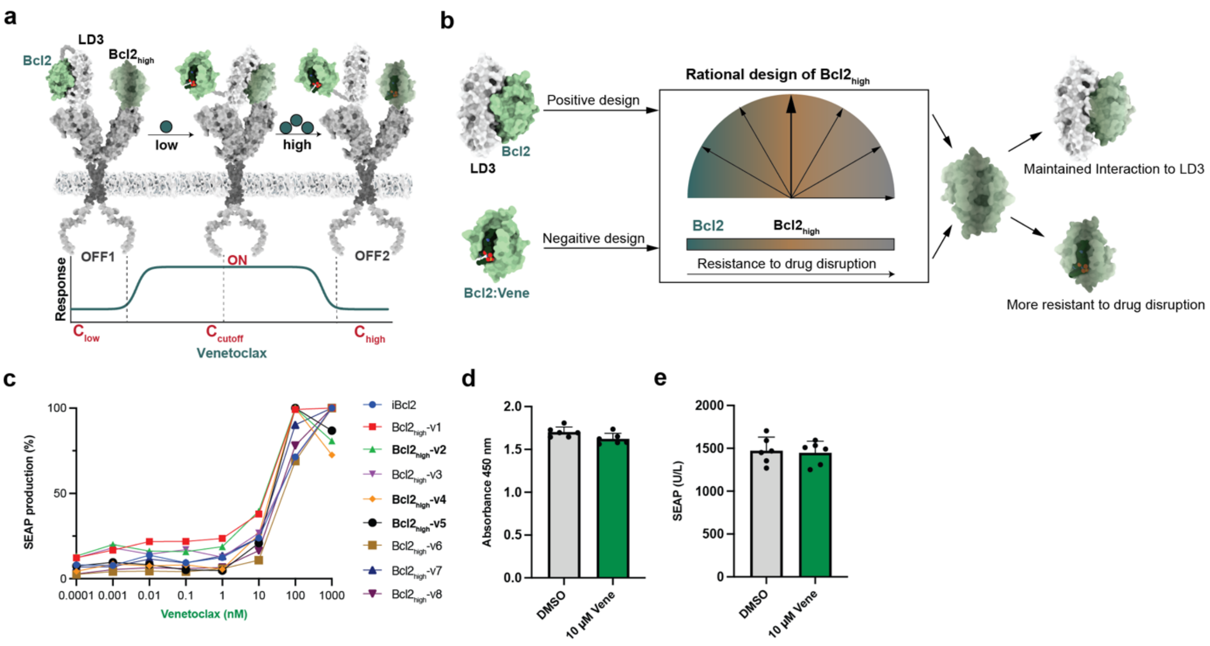


**Supplementary Figure 6: Design, screening and validation of Venetoclax controlled bandpass filters.**

**a)** Bcl2$_{high}$-based bandpass filter designs were developed and to respond to different concentrations of Venetoclax. **b)** Multistate design for tuning drug resistance of drug-receptor protein towards to its corresponding inhibitor. **c)** Screening of Bcl2$_{high}$-based bandpass filter variants in the GEMS platform. The designed binder LD3 was fused to the drug$_{low}$-receptor (PSV40-IgK-Bcl2-GGGGSX3-LD3-EpoRm-IL-6RBm-pA) in one EpoR chain and Bcl2$_{high}$ variants with the insensitive Bcl2 (iBcl2) as control in the other EpoR chain (PSV40-IgK-Bcl2$_{high}$-EpoRm-IL-6RBm-pA). HEK293T cells were transfected with indicated plasmids, A-1155463 drug ranging from 1pM to 1 µM were added 12 hours post-transfection, then SEAP was measured 24 hours after drug treatment. **d-e)** Venetoclax effects on HEK293T cells. HEK293T cells were transfected with a plasmid for constitutive SEAP production and incubated with indicated drugs for 24 hours. SEAP quatification were performed to measure the protein production and WST-8 was used to test the cell proliferation. For cell proliferation assay, WST-8 was added after

24 hours of drug incubation, and measured its absorbance at 450 nm 4 hours later. Each bar represents the mean of six biological replicates ± s.d., overlaid with a scatter dot plot of the original data points.

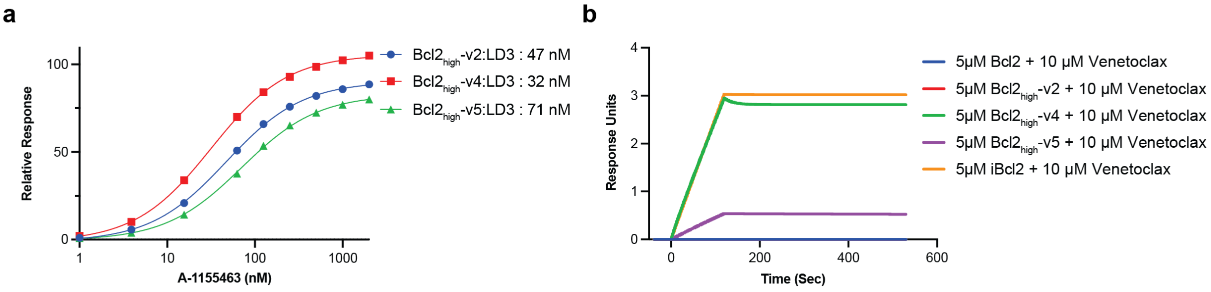


**Supplementary Figure 7: Binding affinities and drug resistances of Bcl2$_{high}$-v(2,4,5) assessed by SPR.**

**a)** Measurement of dissociation constants for Bcl2$_{high}$-v(2,4,5) and LD3 interactions. Equilibrium curves are fitted using three-parameter non-linear regression. **b)** Comparison of A-1155463 resistance of Bcl2$_{high}$-v(2,4,5) measured by SPR. 5 μM of Bcl2 or indicated variants were mixed with 10 μM of Venetoclax and injected over an LD3 immobilized chip to analyse the binding response. iBcl2 remained the highest response (orange), and Bcl2 was completely inhibited (blue). Bcl2$_{high}$-v2 (red), Bcl2$_{high}$-v4 (green), Bcl2$_{high}$-v5 (purple) showed intermediate binding responses in the presence of 10 μM Venetoclax.

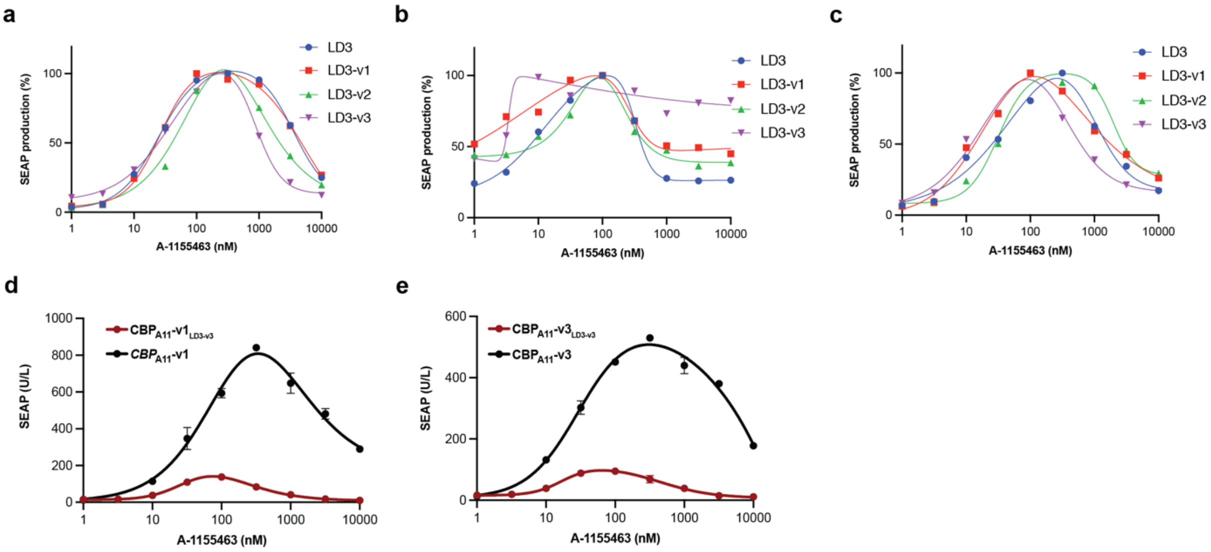


**Supplementary Figure 8: Screening of LD3 variants in CBP$_{A11}$-v(1-3) and raw data of CBP$_{A11}$-v(1,3) with LD3-v3 variant.**

**a-c)** Screening of LD3 and LD3-v(1-3) variants in complex with BclXL for more sensitive CBP$_{A11}$-v1 (a), CBP$_{A11}$-v2 (b), CBP$_{A11}$-v3 (c) in the GEMS platform. The LD3 variants replaced the role of LD3 (PSV40-IgK-BclXL-GGGGSX3-LD3-v(1-3)-EpoRm-IL-6RBm-pA) in one EpoR chain and BclXL$_{high}$-v(1-3) in the other EpoR chain (PSV40-IgK-BclXL$_{high}$-V(1-3)-EpoRm-IL-6RBm-pA). HEK293T cells were transfected with indicated plasmids, A-1155463 drug ranging from 1 pM to 1 μM were added 12 hours post-transfection, then SEAP was measured 24 hours after drug treatment. **d)** Dose responses of CBP$_{A11}$-v1 compared with CBP$_{A11}$-v1$_{LD3\text{-}v3}$ in engineered cells. Each data point represents the mean ±s.d. of three replicates and the curves were calculated using Bell-shaped fitting. **e)** Dose responses of CBP$_{A11}$-v3 compared with CBP$_{A11}$-v3$_{LD3\text{-}v3}$ in engineered cells. Each data point represents the mean ±s.d. of three replicates and the curves were calculated using Bell-shaped fitting.

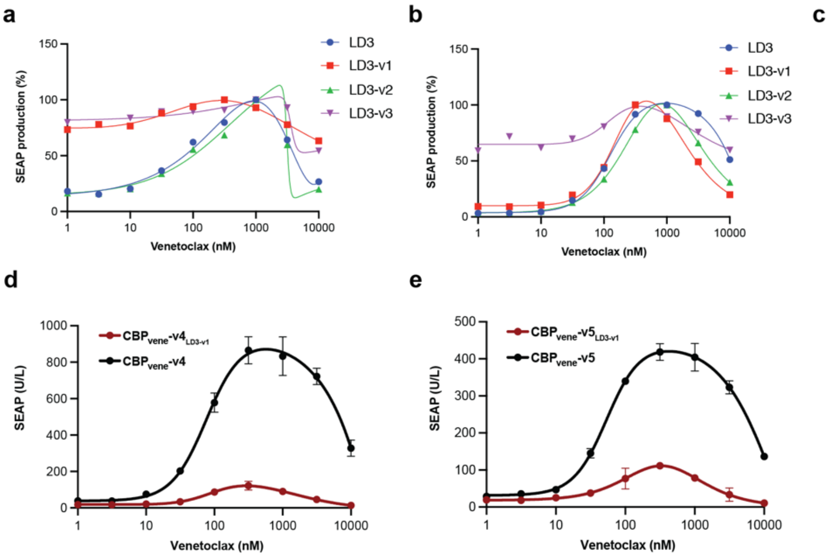


**Supplementary Figure 9: Screening of LD3 variants in CBP$_{vene}$-v(2,4,5) and raw data of CBP$_{vene}$-v(4,5) with LD3-v1.**

**a-c)** Screening of LD3 and LD3-v(1-3) variants in complex with Bcl2 for more sensitive CBP$_{vene}$-v2 (a), CBP$_{vene}$-v4 (b), CBP$_{vene}$-v5 (c) in the GEMS platform. The LD3 variants replaced the role of LD3 (PSV40-IgK-Bcl2-GGGGSX3-LD3-v(1-3)-EpoRm-IL-6RBm-pA) in one EpoR chain and Bcl2$_{high}$-v(2,4,5) in the other EpoR chain (PSV40-IgK-Bcl2$_{high}$-v(2,4,5)-EpoRm-IL-6RBm-pA). HEK293T cells were transfected with indicated plasmids, A-1155463 drug ranging from 1 pM to 1 µM were added 12 hours post-transfection, then SEAP was measured 24 hours after drug treatment. **d)** Dose responses of CBP$_{vene}$-v4 compared with CBP$_{vene}$-v4$_{LD3\text{-}v1}$ in engineered cells. Each data point represents the mean ± s.d. of three replicates and the curves were calculated using Bell-shaped fitting. **e)** Dose responses of CBP$_{vene}$-v5 compared with CBP$_{vene}$-v5$_{LD3\text{-}v1}$ in engineered cells. Each data point represents the mean ± s.d. of three replicates and the curves were calculated using Bell-shaped fitting.

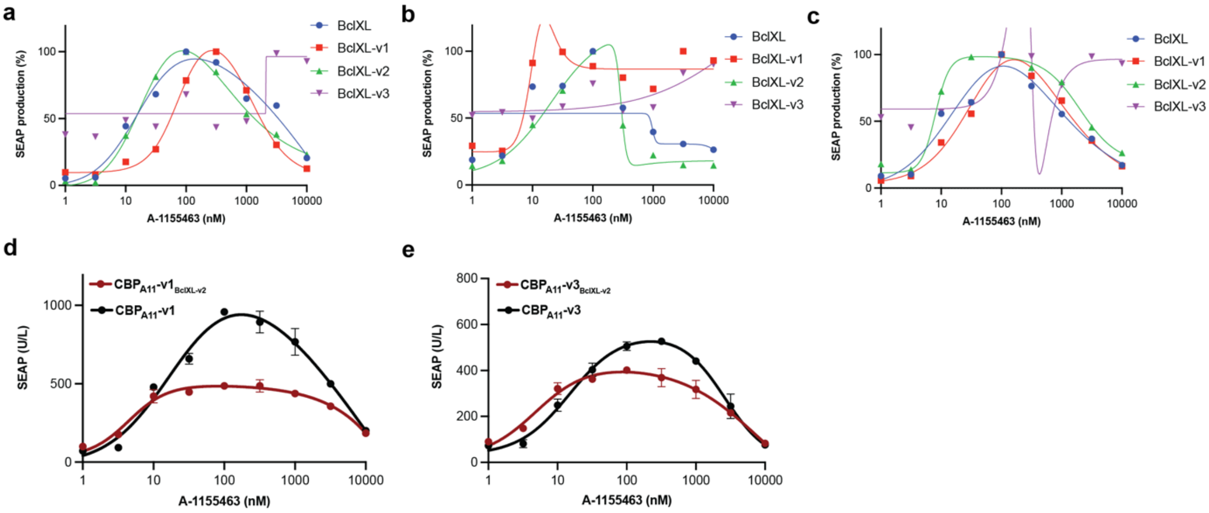


**Supplementary Figure 10: Screening of BclXL Alamut variants in CBP$_{A11}$-v(1-3) and raw data of CBP$_{A11}$-v(1,3) with BclXL-v2.**

**a-c)** Screening of BclXL and BclXL-v(1-3) variants in complex with LD3 for better CBP$_{A11}$-v1 (a), CBP$_{A11}$-v2 (b), CBP$_{A11}$-v3 (c) in the GEMS platform. The BclXL variants replaced the role of LD3 (PSV40-IgK-BclXL-v(1-3)-GGGGSX3-LD3-EpoRm-IL-6RBm-pA) in one EpoR chain and BclXL$_{high}$-v(1-3) in the other EpoR chain (PSV40-IgK-BclXL$_{high}$-v(1-3)-EpoRm-IL-6RBm-pA). HEK293T cells were transfected with indicated plasmids, A-1155463 drug ranging from 1 pM to 1 μM were added 12 hours post-transfection, then SEAP was measured 24 hours after drug treatment. **d)** Drug dose-dependent responses of CBP$_{A11}$-v1 compared with CBP$_{A11}$-v1$_{BclXL\text{-}v2}$ in engineered cells. Each data point represents the mean ±s.d. of three replicates and the curves were calculated using Bell-shaped fitting. **e)** Drug dose-dependent responses of CBP$_{A11}$-v3 compared with CBP$_{A11}$-v3$_{BclXL\text{-}v2}$ in engineered cells. Each data point represents the mean ±s.d. of three replicates and the curves were calculated using Bell-shaped fitting.

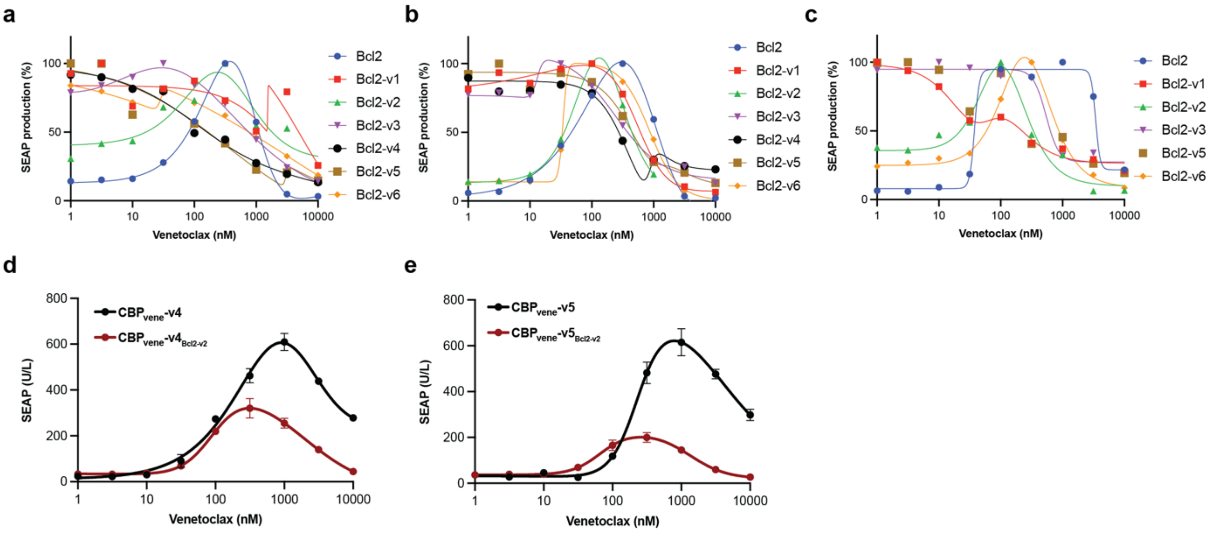


**Supplementary Figure 11: Screening of Bcl2 variants in CBP$_{vene}$-v(2,4,5) and raw data of CBP$_{vene}$-v(2,4,5) with Bcl2-v2.**

**a-c)** Screening of Bcl2 and Bcl2-v(1-6) variants in complex with Bcl2 for more sensitive CBP$_{vene}$-v2 (a), CBP$_{vene}$-v4 (b), CBP$_{vene}$-v5 (c) in the GEMS platform The LD3 variants replaced the role of LD3 (PSV40-IgK-Bcl2-v(1-6)-GGGGSX3-LD3-EpoRm-IL-6RBm-pA) in one EpoR chain and Bcl2$_{high}$-v(2,4,5) in the other EpoR chain (PSV40-IgK-Bcl2$_{high}$-v(2,4,5)-EpoRm-IL-6RBm-pA). HEK293T cells were transfected with indicated plasmids, A-1155463 drug ranging from 1pM to 1 μM were added 12 hours post-transfection, then SEAP was measured 24 hours after drug treatment. **d)** Drug dose-dependent responses of CBP$_{vene}$-v4 compared with CBP$_{vene}$-v4$_{Bcl2\text{-}v2}$ in engineered cells. Each data point represents the mean ±s.d. of three replicates and the curves were calculated using Bell-shaped fitting. **e)** Drug dose-dependent responses of CBP$_{vene}$-v5 compared with CBP$_{vene}$-v5$_{Bcl2\text{-}v2}$ in engineered cells. Each data point represents the mean±s.d. of three replicates and the curves were calculated using Bell-shaped fitting.

### Supplementary Table

**Supplementary Table 1: Single drug reversible design of BclXL.**

| Name | Sequences |
|---|---|
| BclXL$_{high}$-v1 | MSQSNRELVVDFLSYKLSQKGYSWSQFSDVEENRTEAPEGTESEAVKQALREAGD EFELRYRRAFSDLLSQLHITPGTAYQSFEQVVNELFRDGVNWGRIVAFFSFGGLLCV ESVDKEMQVLVSRIAAWMATYLNDHLEPWIQENGGWDTFVELYGNNAAAESRKGQ ER |
| BclXL$_{high}$-v2 | MSQSNRELVVDFLSYKLSQKGYSWSQFSDVEENRTEAPEGTESEAVKQALREAGD EFELRYRRAFSDLTSQLHITPGTAYQSFEQVVNELFRDGVNWGRIVAFFSFGGVLC VESVDKEMQVLVSRIAAWMATYLNDHLEPWIQENGGWDTFVELYGNNAAAESRKG QER |
| BclXL$_{high}$-v3 | MSQSNRELVVDFLSYKLSQKGYSWSQFSDVEENRTEAPEGTESEAVKQALREAGD EFELRYFRAFSDLVSQLHITPGTAYQSFEQVVNELFRDGVNWGRIVAFFSFGGALCV ESVDKEMQVLVSRIAAWMATYLNDHLEPWIQENGGWDTFVELYGNNAAAESRKGQ ER |
| BclXL$_{high}$-v4 | MSQSNRELVVDFLSYKLSQKGYSWSQFSDVEENRTEAPEGTESEAVKQALREAGD EFSLRYERAISDLTSQLHITPGTAYQSFEQVVNELFRDGVNWGRIVAFFSFGGALCV ESVDKEMQVLVSRIAAWMATYLNDHLEPWIQENGGWDTFVELYGNNAAAESRKGQ ER |
| BclXL$_{high}$-v5 | MSQSNRELVVDFLSYKLSQKGYSWSQFSDVEENRTEAPEGTESEAVKQALREAGD EFELRYERAISDLVSQLHITPGTAYQSFEQVVNELFRDGVNWGRIVAFFSFGGALCV ESVDKEMQVLVSRIAAWMATYLNDHLEPWIQENGGWDTFVELYGNNAAAESRKGQ ER |
| BclXL$_{high}$-v6 | MSQSNRELVVDFLSYKLSQKGYSWSQFSDVEENRTEAPEGTESEAVKQALREAGD EFSLRYERAISDLVSQLHITPGTAYQSFEQVVNELFRDGVNWGRIVAFFSFGGALCV ESVDKEMQVLVSRIAAWMATYLNDHLEPWIQENGGWDTFVELYGNNAAAESRKGQ ER |

**Supplementary Table 2: Single drug reversible Bcl2.**

| Name | Sequences |
|---|---|
| Bcl2$_{high}$-v1 | MAHPGRTGYDNREIVMKYIHYKLSQRGYEWDAGDDVEENRTEAPEGTESEVVHLTLR QAGDDFSRRYRRDFAEMSSQLHLTPFTARGRFATVVEELFRDGVNWGRIVAFFEFGG VMCIESVNREMSPLVDNIALWMTEYLNRHLHTWIQDNGGWDAFVELHGPSMR |
| Bcl2$_{high}$-v2 | MAHPGRTGYDNREIVMKYIHYKLSQRGYEWDAGDDVEENRTEAPEGTESEVVHLTLR QAGDNFSRRYRRDFAEMSSQLHLTPFTARGRFATVVEELFRDGVNWGRIVAFFEFGG VMCVESVNREMSPLVDNIALWMTEYLNRHLHTWIQDNGGWDAFVELHGPSMR |
| Bcl2$_{high}$-v3 | MAHPGRTGYDNREIVMKYIHYKLSQRGYEWDAGDDVEENRTEAPEGTESEVVHLTLR QTGDSFSRRYRRDFAEMSSQLHLTPFTARGRFATVVEELFRDGVNWGRIVAFFEFGG VMCVESVNREMSPLVDNIALWMTEYLNRHLHTWIQDNGGWDAFVELYGPSMR |
| Bcl2$_{high}$-v4 | MAHPGRTGYDNREIVMKYIHYKLSQRGYEWDAGDDVEENRTEAPEGTESEVVHLTLR QAVDDFSRRYRRDFAEMSSQLHLTPFTARGRFATVVEELFRDGVNWGRIVAFFEFGG VMCVESVNREMSPLVDNIALWMTEYLNRHLHTWIQDNGGWDAFVELYGPSMR |

| | |
|---|---|
| Bcl2$_{high}$-v5 | MAHPGRTGYDNREIVMKYIHYKLSQRGYEWDAGDDVEENRTEAPEGTESEVVHLTLR<br>QAGD**Y**FSRRYRRDFAEMSSQLHLTPFTARGRFATVVEELFRDGVNWGRIVAFFEFGG<br>VMCVESVNREMSPLVDNIALWMTEYLNRHLHTWIQDNGGWDAFVELYGPSMR |
| Bcl2$_{high}$-v6 | MAHPGRTGYDNREIVMKYIHYKLSQRGYEWDAGDDVEENRTEAPEGTESEVVHLTLR<br>QA**V**D**Y**FSRRYRRDFAEMSSQLHLTPFTARGRFATVVEELFRDGVNWGRIVAFFEFGG<br>VMCVESVNREMSPLVDNIALWMTEYLNRHLHTWIQDNGGWDAFVELYGPSMR |
| Bcl2$_{high}$-v7 | MAHPGRTGYDNREIVMKYIHYKLSQRGYEWDAGDDVEENRTEAPEGTESEVVHLTLR<br>QAGD**E**FSRRYRRDFAEMSSQLHLTPFTARGRFATVVEELFRDGVNWGRIVAFFEFGG<br>VMCVESVNREMSPLVDNIALWMTEYLNRHLHTWIQDNGGWDAFVELYGPSMR |
| Bcl2$_{high}$-v8 | MAHPGRTGYDNREIVMKYIHYKLSQRGYEWDAGDDVEENRTEAPEGTESEVVHLTLR<br>QA**V**D**E**FSRRYRRDFAEMSSQLHLTPFTARGRFATVVEELFRDGVNWGRIVAFFEFGG<br>VMCVESVNREMSPLVDNIALWMTEYLNRHLHTWIQDNGGWDAFVELYGPSMR |

**Supplementary Table 3: LD3 variants.**

| Name | Sequences |
|---|---|
| LD3-v1 | QRWELALGRFL**A**YLSWVSTLSEQVQEELLSSQVTQELRALMDETMKELKAYKSELEEQL<br>TPVAEETRARLSKELQAAQARLGADMEDVRGRLVQYRGEVQAMLGQSTEELRVRLASH<br>LIALALRLIGDAFDLQKRLAVY |
| LD3-v2 | QRWELALGRFLEYLSWVSTLSEQVQEELLSSQVTQELRALMDETMKELKAYKSELEEQL<br>TPVAEETRARLSKELQAAQARLGADMEDVRGRLVQYRGEVQAMLGQSTEELRVRLASH<br>LIALAL**A**LIGDAFDLQKRLAVY |
| LD3-v3 | QRWELALGRFLEYLSWVSTLSEQVQEELLSSQVTQELRALMDETMKELKAYKSELEEQL<br>TPVAEETRARLSKELQAAQARLGADMEDVRGRLVQYRGEVQAMLGQSTEELRVRLASH<br>LIALALRLIG**A**AFDLQKRLAVY |

**Supplementary Table 4: BclXL variants.**

| Name | Sequences |
|---|---|
| BclXL-v1 | MSQSNRELVVDFLSYKLSQKGYSWSQFSDVEENRTEAPEGTESEAVKQALREAGDEF<br>ELRYRRAFSDLTS**A**LHITPGTAYQSFEQVVNELFRDGVNWGRIVAFFSFGGALCVESV<br>DKEMQVLVSRIAAWMATYLNDHLEPWIQENGGWDTFVELYGNNAAAESRKGQER |
| BclXL-v2 | MSQSNRELVVDFLSYKLSQKGYSWSQFSDVEENRTEAPEGTESEAVKQALREAGDEF<br>ELRYRRAFSDLTSQLHITPGTAYQSFEQVVNE**A**FRDGVNWGRIVAFFSFGGALCVESV<br>DKEMQVLVSRIAAWMATYLNDHLEPWIQENGGWDTFVELYGNNAAAESRKGQER |
| BclXL-v3 | MSQSNRELVVDFLSYKLSQKGYSWSQFSDVEENRTEAPEGTESEAVKQALREAGDEF<br>ELRYRRAFSDLTSQLHITPGTAYQSFEQVVNELFRDGVNWG**A**IVAFFSFGGALCVESV<br>DKEMQVLVSRIAAWMATYLNDHLEPWIQENGGWDTFVELYGNNAAAESRKGQER |

**Supplementary Table 5: Bcl2 variants.**

| Name | Sequences |
|---|---|
| Bcl2-v1 | MAHPGRTGYDNREIVMKYIHYKLSQRGYEWDAGDDVEENRTEAPEGTESEVVHLTLR<br>QAGDD**A**SRRYRRDFAEMSSQLHLTPFTARGRFATVVEELFRDGVNWGRIVAFFEFGG<br>VMCVESVNREMSPLVDNIALWMTEYLNRHLHTWIQDNGGWDAFVELYGPSMR |
| Bcl2-v2 | MAHPGRTGYDNREIVMKYIHYKLSQRGYEWDAGDDVEENRTEAPEGTESEVVHLTLR<br>QAGDDFSRRYRRDFAEMSSQLHLTPFTARGRFAT**A**VEELFRDGVNWGRIVAFFEFGG<br>VMCVESVNREMSPLVDNIALWMTEYLNRHLHTWIQDNGGWDAFVELYGPSMR |
| Bcl2-v3 | MAHPGRTGYDNREIVMKYIHYKLSQRGYEWDAGDDVEENRTEAPEGTESEVVHLTLR<br>QAGDDFSRRYRRDFAEMSSQLHLTPFTARGRFATVVE**A**LFRDGVNWGRIVAFFEFGG<br>VMCVESVNREMSPLVDNIALWMTEYLNRHLHTWIQDNGGWDAFVELYGPSMR |
| Bcl2-v4 | MAHPGRTGYDNREIVMKYIHYKLSQRGYEWDAGDDVEENRTEAPEGTESEVVHLTLR<br>QAGDDFSRRYRRDFAEMSSQLHLTPFTARGRFATVVEE**A**FRDGVNWGRIVAFFEFGG<br>VMCVESVNREMSPLVDNIALWMTEYLNRHLHTWIQDNGGWDAFVELYGPSMR |
| Bcl2-v5 | MAHPGRTGYDNREIVMKYIHYKLSQRGYEWDAGDDVEENRTEAPEGTESEVVHLTLR<br>QAGDDFSRRYRRDFAEMSSQLHLTPFTARGRFATVVEELFRDGVNWG**A**IVAFFEFGG<br>VMCVESVNREMSPLVDNIALWMTEYLNRHLHTWIQDNGGWDAFVELYGPSMR |
| Bcl2-v6 | MAHPGRTGYDNREIVMKYIHYKLSQRGYEWDAGDDVEENRTEAPEGTESEVVHLTLR<br>QAGDDFSRRYRRDFAEMSSQLHLTPFTARGRFATVVEELFRDGVNWGRIVAFF**A**FGG<br>VMCVESVNREMSPLVDNIALWMTEYLNRHLHTWIQDNGGWDAFVELYGPSMR |
